# The Evolutionary Landscape of Pan-Cancer Drives Clinical Aggression

**DOI:** 10.1101/422667

**Authors:** Shichao Pang, Leilei Wu, Xin Shen, Yidi Sun, Jingfang Wang, Yi-Lei Zhao, Zhen Wang, Yixue Li

**Author notes:** Correspondence and requests for materials should be addressed to Y.X.L., Z.W., Y.-L.Z. or J.F.W.

## Abstract

Although cancer mechanisms differ from occurrence and development, some of them have similar oncogenesis, which leads to similar clinical phenotypes. Most existing genotyping studies look at “omics” data, but intentionally or unintentionally avoided that cancer is a time-dependent evolutionary process, biologically represented by the time evolution of tumor clones. We used the Bayesian mutation landscape approach to reconstruct the evolutionary process of cancer by acquiring somatic mutation data consisting of 21 cancer types. Four representative evolution patterns of pan-cancer have been discovered: trees, chaos, biconvex, and Cambrian, and a strong correlation between these four evolutionary patterns and clinical aggressivity. We further explained the characteristics of the corresponding biological systems in the evolution of pan cancer by analyzing the function of differentially expressed protein-protein interaction networks. Our results explained the difference in clinical aggressivity between cancer evolution patterns from the evolution of tumor clones and exposed the functional mechanism behind.

## Introduction

Cancer is a multistage process that abnormal cells invade or spread to other parts of the body(Plummer et al. 2016), causing about 15.7% of human deaths(Wang et al. 2016). Different cancers vary a lot in prognosis and exacerbation. For example, patients with breast tumor have a 72% 5-year survival rate in stage III, but only 3% pancreatic patients can survive after 5 years(Howlader N, Noone AM, Krapcho M, Miller D, Bishop K, Kosary CL, Yu M, Ruhl J, Tatalovich Z, Mariotto A, Lewis DR, Chen HS, Feuer EJ 1975). Usually, similar oncogenesis will lead to similar clinical outcomes. For instance, different type of cancers sometimes positively respond to the same chemical analogous and vaccine(Howell-Jones et al. 2010), and share similar mutation frequency of genes in background for the related opening area and frequency of the double helix DNA strands(Perry Evans, Stefan Avey, Yong Kong 2013). This is the starting point of pan-cancer researches. Scientists have tried diverse methods to identify pan-cancer pattern using omics data, e.g., somatic nucleotide variants (SNV)(Leiserson et al. 2015), copy number variation (CNV)(Zack et al. 2013), proteomics(Zhang et al. 2014) and DNA methylation(Yang et al. 2017). But the results are not as expected, because the occurrence and development of cancers is a time-dependent evolutionary process. Recent studies indicated that the tumor aggressivity always links to its heterogeneity(Jögi et al. 2012), and reflects in clinical outcomes. Analysis of cancer evolutionary process combined with time-dependent survivals could help us to figure out the clinical aggressivity of tumors.

Cancers can be viewed as an evolutionary process based on the clonal selection and dynamic process of immune responses(Gong et al. 2009). The accumulation of somatic mutations during clonal expansion, combined with microenvironment variations(Nowell and Nowell PC. 1976), drives the evolutionary changes of tumor cells. The stochastic process is the theoretical foundation of cancer evolution. For instance, the linear theory came out in 2003(Nowak et al. 2003) compared the cancer evolution process with the Moran process(Nowak et al. 2003). Following nonlinear and branching theory(Anderson et al. 2011) reminded us to pay more attention to subclones and explore possible paths for cancer progression. In 2015, the big bang theory raised the idea that tumor expanded predominantly from an early clone mixed with numerous subclones(Sottoriva et al. 2015). Besides, recent studies also put forward a neutral evolutionary theory(Williams et al. 2016), similar to Kimura’s(Kimura 1977). Our previous study on clear cell renal carcinoma reconstructed a phylogenetic tree model in a fashion of stage-by-stage expansion(Pang et al. 2018). Since these theories were based on the studies of different cancers, we need to use a uniform algorithm to figure out the evolution patterns of pan-cancer.

In the current study, we reconstructed the evolution processes for pan-cancers with somatic mutations across pathological stages, based on which four representative evolutionary patterns (tree, chaos, biconvex and Cambrian) were proposed. Then we analyzed the similarities and differences of clinical aggressivity for these evolutionary patterns. We further explained the functional characteristics of the pan-cancer evolution pattern by a protein-protein interaction network based on the differentially expressed genes.

## Results

### Mutation and survival landscape of pan-cancer

We collected clinical and genomic data sets of 21 types of cancers from the Cancer Genome Atlas (TCGA) cohort (The full names of cancers were listed in **Table S1**). Since not all of them were well-paired, we finally chose 5,134 samples with somatic SNVs for constructing evolution processes and 9,249 samples for survival analysis. The annotation information on related biological system, early detection of cancer, tumor type and M/C class were showed in **Table S1**.

The gene mutation landscape indicated that mutation frequency differed among different types of cancers. For instance, SKCM and UCEC had high discreteness, while KIRC and THCA were centralized (**Fig. 1a**). Additionally, gene mutation frequency did not increase with the progress of pathological stages in most cancers (**Fig. 1b, Table S2**). Although mutation frequency always correlated with tumor deterioration for specific cancers, general survival outcome didn’t exhibit a consistence among pan-cancer (**Fig. 1c**). For example, even with a relatively low mutation frequency, OV showed a poor 5-year survival rate. Then we carried out a hierarchical cluster analysis with a combination of mutation frequency and 5-year survival rate (**Fig. 1d**). In the yellow box cluster, both OV and LUSC showed poor survival outcomes, but LUSC possessed a high mutation frequency. As survival rate is a time corresponding symptom of cancers, we reconstructed evolution processes for cancers across pathological stages to figure out the similarities and differences of oncogenesis in pancancer.

**Figure 1:**
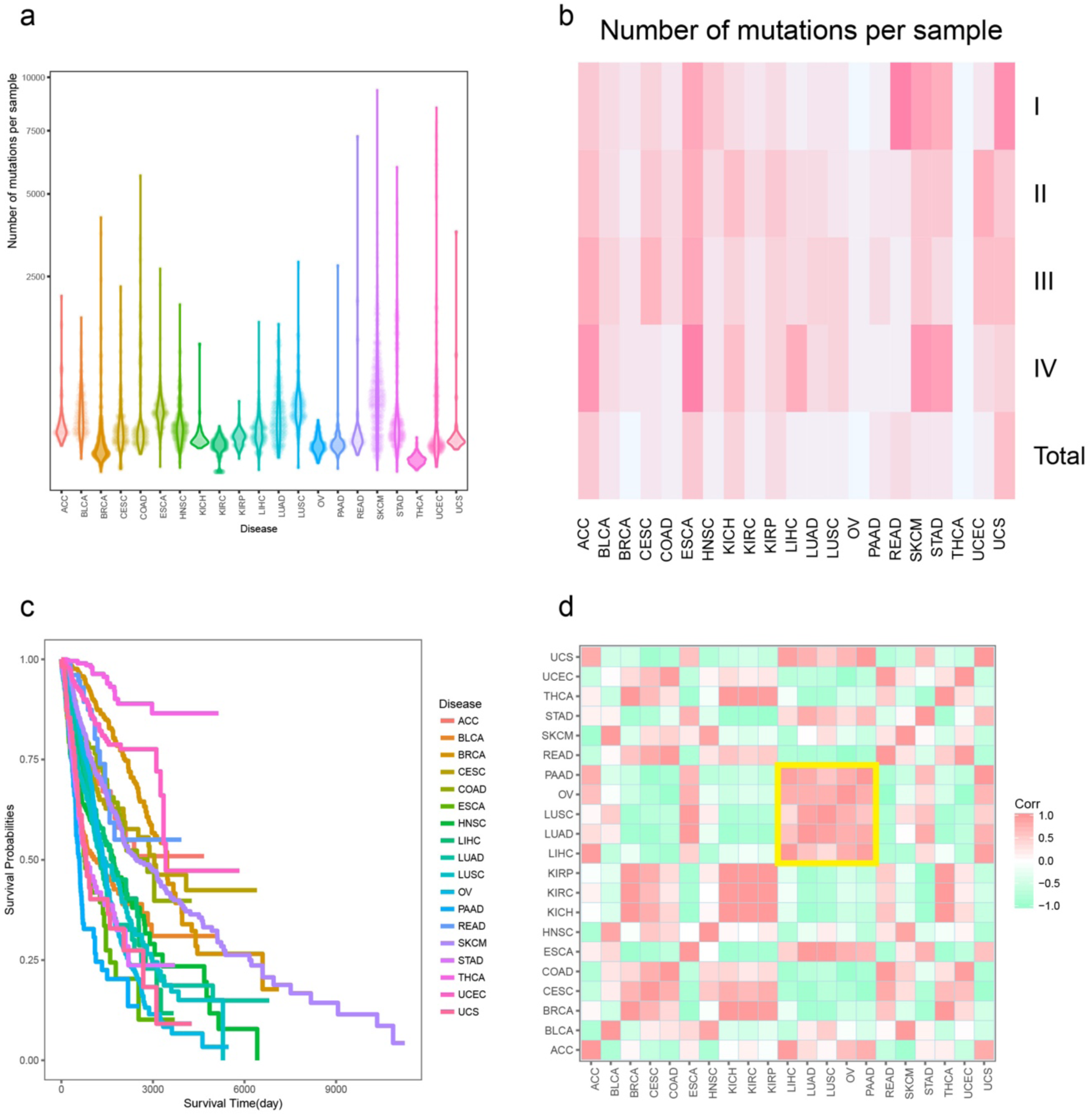
Mutation and clinical landscape of 21 types of cancers. (A). Mutation frequency of 21 types of cancers. (B). Mutation frequency of 21 types of cancers in each pathological stages. (C). Survival curve of 21 types of cancers (Kaplan-Meier estimator). (D). Correlation heatmap of mutation (median mutation frequency) and survival (5-year survival rate) features in 21 types of cancers.

### Reconstruction of pan-cancer evolution process and NMF cluster-based pattern

Since genetic studies always focused on high-frequency mutations, the evolutionary path and essential variations with moderate frequency were missing generally. We employed the Bayesian Mutation Landscape (BML) methods to reconstruct evolutionary processes based on somatic mutations, and generated directed acyclic graphs (DAGs) of each cancer using four pathological stages representing four-time points during the tumor progression (**Fig. S1**). The bootstrap method was used to extract information with a highly statistical confidence (for detailed information, please see **Methods**). A total of 12 features were extracted, including DAG nodes, edges and key genes in each pathological stage. Here, we defined key genes as those appearing in more than one pathological stage. Interestingly, the four vectors extracted for nonnegative matrix factorization (NMF) clusters coincide with pathological stages: vector 3 and vector 4 were mainly contributed by stage I and II, respectively; vector 2 and vector 1 were mainly contributed by stage III and IV. In addition, stage III also had a slight contribution to vector 1 (**Fig. S2**). Finally, we generated four evolution patterns for cancers based on the NMF clusters (**Fig. 2a, 2b and 2c**).

**Figure 2:**
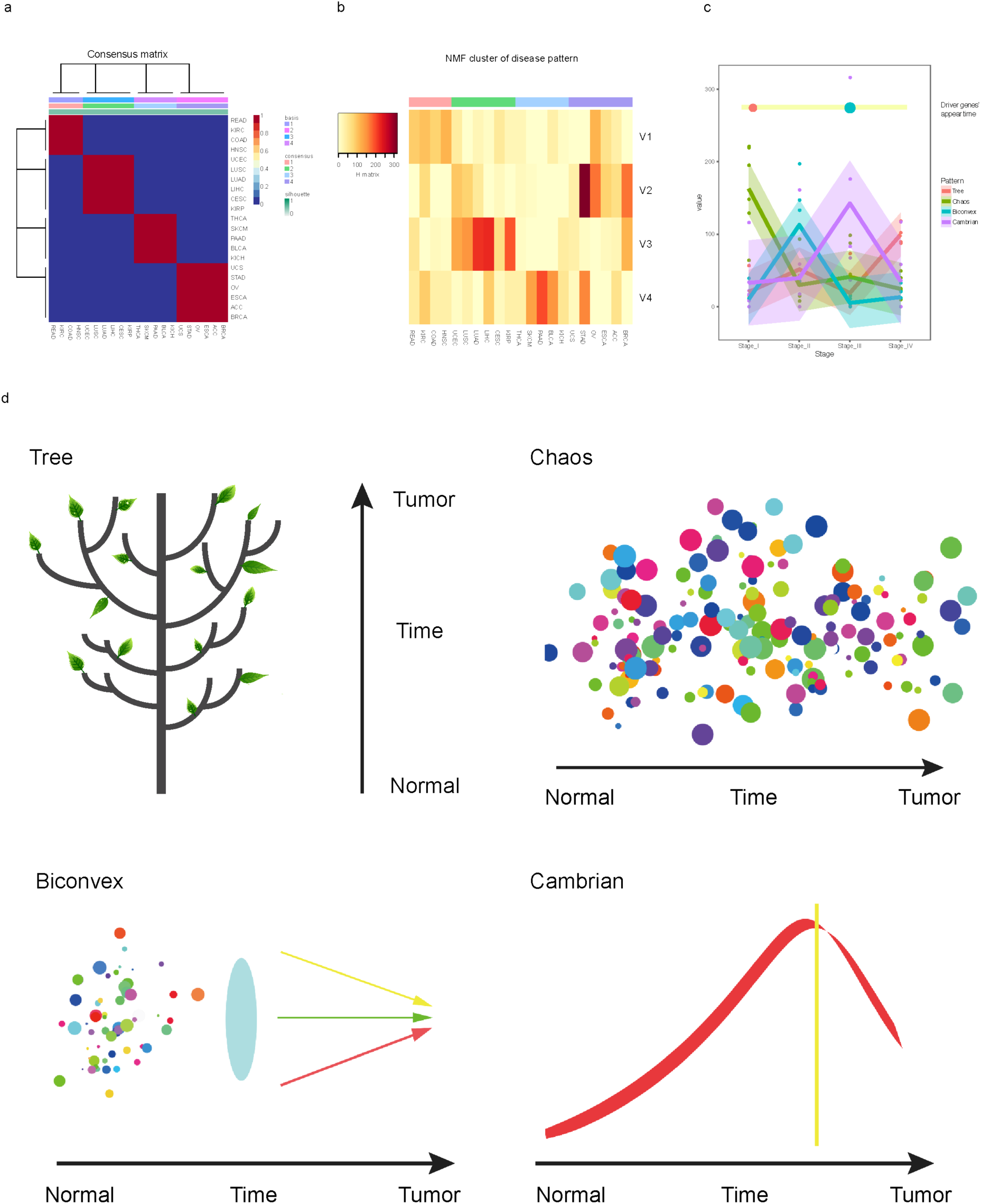
Evolution pattern of 21 types of cancers. (A). Consensus map of pan-cancer NMF cluster. Basis represented four vectors in Figure3b and consensus represented four clusters. (B). Coefficient map of pan-cancer NMF cluster. (C). Pan-cancer evolution process across stages according to NMF cluster result. (D). Schematic diagram of four cancer evolution patterns from normal to tumor. Tree: High order evolution process with dominant driver genes. Chaos: No dominant driver genes and multiple kinds of evolutionary paths. Biconvex: Joint of early-chaos and late-tree, and had dominant driver genes in late stage. Cambrian: Peaceful early stage combined with explosion of gene mutation and evolutionary paths in late stage.

The first cluster had no significantly dominated vector, and only vector 1 showed a slight advantage. KIRC and READ had three vectors with remarkable mixture coefficient (H matrix) while HNSC and COAD had two. Tumors in this pattern showed major evolutionary paths in DAGs, and progressed smoothly. Driver genes with high-frequency mutations (e.g., VHL gene in KIRC and APC gene in COAD) appeared in early pathological stages, and were close to normal node in DAGs of all pathological stages. This process is similar to the growth of trees, so-named “tree” pattern. From the perspective of competitive evolution of tumor cloning, the “tree” model indicates that certain tumor clones dominate tumorigenesis and development, and tumor clones presents a competitive equilibrium with each other, and tumor heterogeneity is low in this case. The second cluster was dominated by vector 3, while mixture coefficient of other vectors in this cluster were in average. No driver genes were found in DAGs of this cluster. Instead of major evolutionary paths, tumors in this cluster like CESC exhibited multiple kinds of evolutionary paths, resulting in highly heterogeneity. Thus, we named it as “chaos” pattern. Unlike the “tree” model, the evolutionary behavior of tumor clones corresponding to the “chaos” pattern presents a competitive evolution caused by the dissemination of a large number of non-dominant clones, and a random equilibrium state that they reach each other, and tumor heterogeneity is high in this case. The third cluster is remarkably dominated by vector 4. Different from the other clusters, vector 3 in this cluster showed a comparatively low mixture coefficient. Limited evolutionary paths were observed in stage I, but more appeared in stage II. Although multiple evolutionary paths appeared in this stage, no one exhibited dominance. In late-stage (III and IV), the mutation frequency of driver genes (e.g., PIK3CA gene in BLCA) increased, and major evolutionary paths were formed. The late-stage performance of this pattern is more smooth due to the appearance of major evolutionary paths. Just like a biconvex to make dispersed light converged, we named this cluster as a “biconvex” pattern. The “biconvex” pattern reflects the different evolutionary patterns of tumor cloning. At the beginning, there are only a small number of tumor clones, and the competitive evolution is in an equilibrium state with no dominant clones. Then, the tumor clone containing the driving gene appears, and through competitive evolution suppresses the survival of other tumor clones and evolves into a dominant tumor clone, and tumor heterogeneity at the final stage is low in this “biconvex” pattern. The fourth cluster is dominated by vector 2 while vector 1 also showed a remarkable mixture coefficient. Tumors in this cluster had moderate number of evolutionary paths and few genes with high-frequency mutations in early-stage. Enormous evolutionary paths spring up since stage III in cancers like BRCA, looking like the Cambrian. So, we called it “Cambrian” pattern. Tumors in this cluster usually had no SNV driver genes, and was not SNV dominated (**Table S1**). The “Cambrian” pattern seems to be exactly the opposite of “biconvex”. At the beginning, only a moderate number of tumor clones occurred, and then in the middle and late stages of tumorigenesis and development, the number of tumor clones suddenly exploded. At this time, the unique pattern of tumor cloning evolution of “chaos” pattern appeared, and a large number of non-dominant tumor clones reached the final competitive evolutionary balance. At the final stage, the tumor exhibits a high degree of heterogeneity.

### Survival outcomes of pan-cancer evolution patterns

After identifying the cancer evolution patterns, we explored survival outcomes for each evolution pattern. Among all the evolution patterns (**Fig. 3a and Table S3**), Cambrian pattern showed a significant distinction in survival outcome between early and late stages. Because increased evolutionary paths in late stages hastened tumor progression, leading to high tumor heterogeneity and causing bad survival outcome. In biconvex pattern, a better survival outcome was found in stage III rather than stage II (Wald test, p-value=0.038, HR: 3.189(2.691∼3.793)). Because scattered evolutionary paths in stage II became disciplinary to form major evolutionary paths in stage III, resulting in decrement of tumor heterogeneity. Chaos and tree patterns had similar survival pattern across different pathological stages. Their survival curves were regular, and the differences between adjacent pathological stages were uniform. As more evolutionary paths lead to high heterogeneity and result in aggressive clinical outcome, tree pattern showed a better survival outcome than chaos pattern. The stage by stage progression is accordant with tree pattern but unexpected for chaos pattern. One possible explanation is that the multiple kinds of evolutionary paths observed in stage I in chaos pattern expanded to tree pattern subclones. The diversity of chaos pattern evolutionary paths and lack of major evolutionary path contributed to its high heterogeneity.

**Figure 3:**
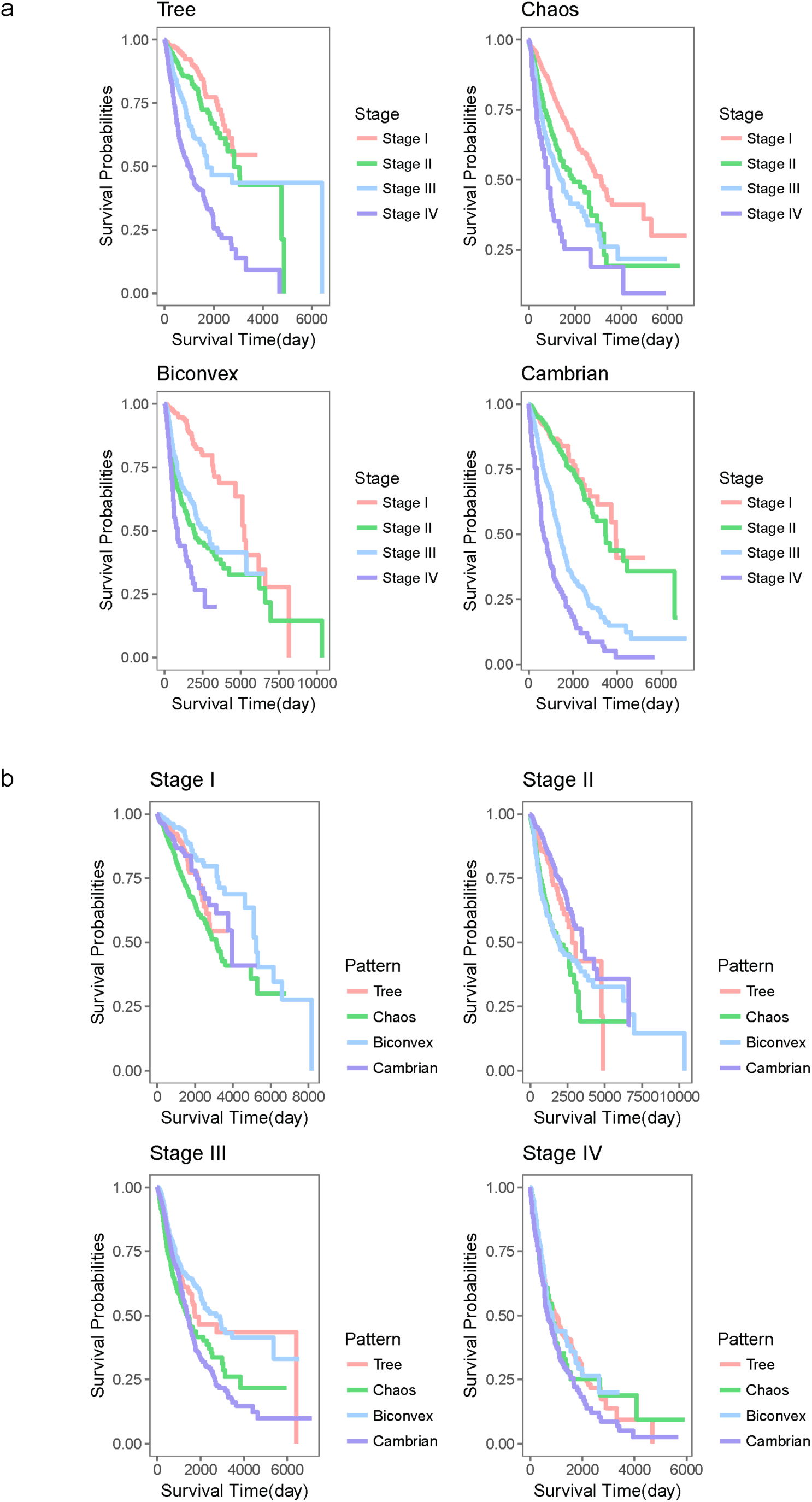
Survival analysis of pan-cancer evolution pattern. (A). Survival outcome of each pathological stages in different evolution pattern. (B). Survival outcome of each cancer evolution pattern in different pathological stages.

Additionally, we also compared the survival outcomes among all the evolution patterns in each pathological stage (**Fig. 3b** and **Table S3**). Cambrian pattern showed a comparatively good survival outcome in early stages. However, its survival outcome turned to be the worst among all the evolution patterns in the last stages. Due to its orderliness, tree pattern exhibited moderate survival outcomes in all pathological stages compared to other evolution patterns. Biconvex pattern had a comparatively lousy survival outcome in stage II due to the similar environment with chaos pattern. After major evolutionary paths formed, the survival curve of biconvex pattern showed high similarity with tree pattern in stage IV (Wald test, p-value=0.97, HR: 0.996(0.777∼1.276)). As expected, chaos pattern employed the worst survival outcomes in almost all pathological stages due to the high tumor heterogeneity.

### Biological function analysis for pan-cancer evolution patterns

We also performed functional analysis for the evolution patterns based on the differentially expressed genes in cancers. We used a threshold of p-value<0.01 and fold-change >3 to detect differentially expressed genes (DEGs) in genomic data of cancers. For each evolution pattern, we merged all tumors and DEGs into a single network based on their belonging relationship (**Fig. 4a**). We also added links between genes according to Human Protein Reference Database (HPRD) protein-protein interactions (PPI). Chaos pattern has the highest network heterogeneity and tree pattern has the most centralized network structure (**Table S4, Fig. S3**). Statistical information for cancer connection degree and PPI degree of each DEGs was represented in **Table S5**. Some DEGs were highly connected to cancers, but their PPI degrees were comparatively low (e.g., FOXM1 and PDK4). They were likely to be a consequence rather than an inducement. However, DEGs with high degrees in both PPI and cancer-connection should be valued. Among these high degree genes, MMP9, MMP2, DES, DCN, COL1A1, SPP1 and CAV1 enriched together in multi-pathways. They functioned together in four cancer evolution pattern.

**Figure 4:**
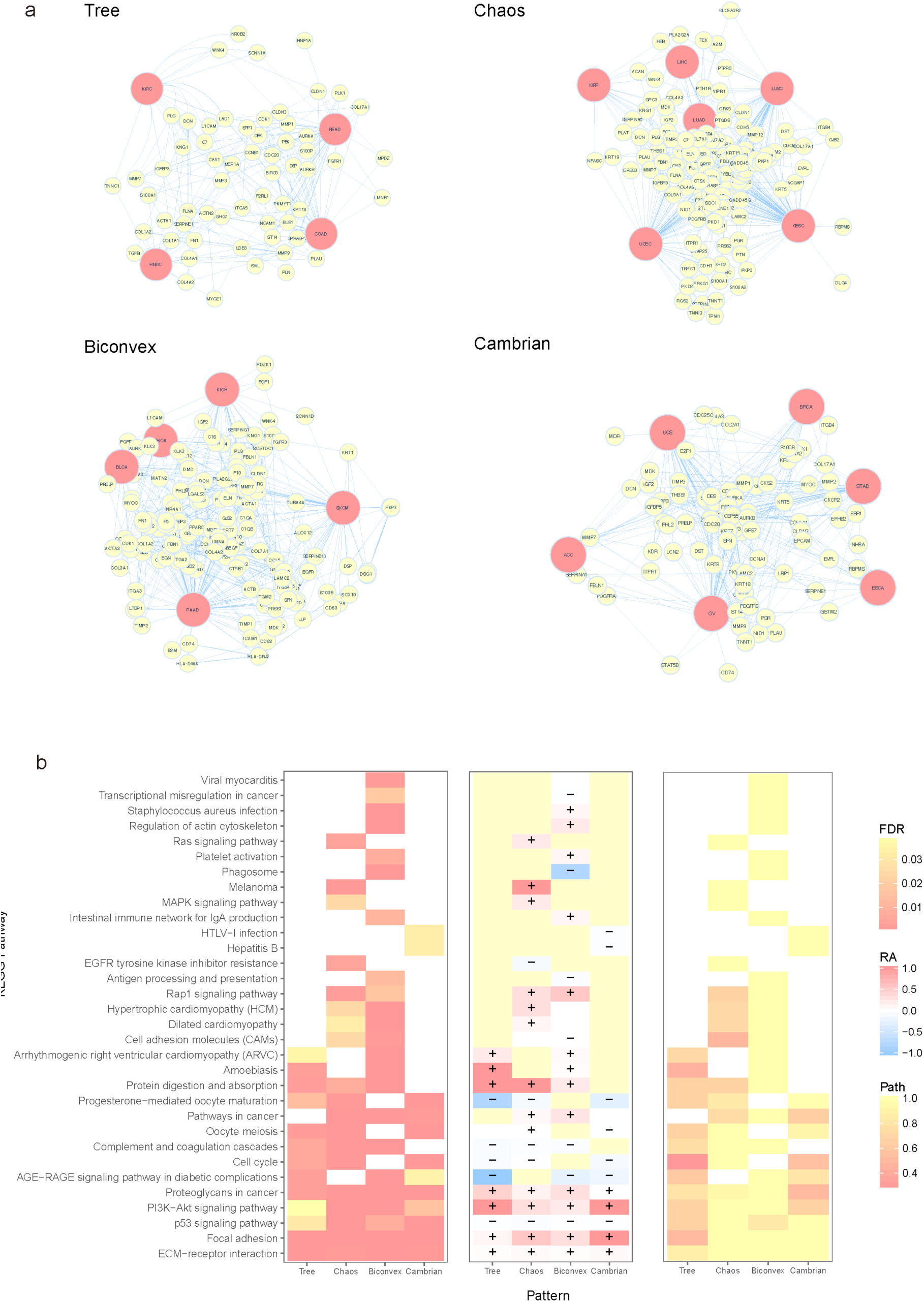
Function analysis of pan-cancer evolution patterns. (A). PPI network of high degree(>5) DEG nodes. (B). KEGG pathway enrichment of high degree nodes (>5) in each cancer evolution pattern PPI network. Enrichment FDR p-value (left), regulation area (middle), paths (right).

Based on the PPI network for differentially expressed genes for each evolution pattern, we picked out high degree DEGs (>=5) for further functional enrichment analysis. Diverse cancer evolution progression needs identical variations of pathways and genes, which always influence the basic functions of cancers. Functional analysis indicated that the evolution patterns shared five biological pathways, i.e., ECM-receptor interaction, focal adhesion, p53 signaling pathway, PI3K-Akt signaling pathway and proteoglycans in cancer (**Fig. 4b**). Although most pattern-shared pathways were confirmed cancer hallmarks, four evolution patterns had their particular pathways, for example, Hepatitis B pathway in Cambrian pattern and MAPK signaling pathway in chaos pattern. Besides, there were seven pathways shared by three evolution patterns, e.g., AGE-RAGE signaling pathway and protein digestion and absorption. They were not often discussed in cancer studies before. But they were closed to inflammation which is a preprocess of cancer(Riehl et al. 2009). AGE-RAGE signaling pathway was absent only in chaos pattern, and functions to increase oxidative stress generation and evoke inflammatory, fibrotic, proliferative, etc. Tree pattern had the least unique pathways, while the unique pathways for chaos and biconvex patterns were more variable, due to their heterogeneity in the early pathological stages. Cambrian pattern didn’t show a lot of exclusive pathways due to its diversity in late pathological stages.

We also evaluated these enriched pathways by DEG locations and directed paths in the same KEGG pathways. Five common pathways showed high similarity in DEG locations. Tree pattern had the least directed paths and highest centralization regulation area, indicating throughout major evolution paths. Despite various evolution paths appeared in Cambrian pattern in the late stages, their functional variation focused on minimum pathways. Most of the pathways were in downstream regulations and directed paths inside pathway were also limited. The explosion seemed to be an effect of system disorders accumulation. Chaos and biconvex patterns showed high similarity in **Fig. 4b**, and had more enriched pathways than the others. Biconvex pattern is consisting of early-chaos and late-tree, which coincident with its survival outcome. Compared to chaos pattern, it had more directed paths and downstream genes. The downstream early-chaos relieved system deterioration and resulted in better survival outcome.

## Discussion

Many investigations have used genome information (e.g., SNV, CNV, and DNA methylation) and proteomic data to perform pan-cancer studies. However, cancer is a time-dependent evolution process and survival outcome is also a time-corresponding symptom. The reconstruction of evolutionary paths is able to provide a novel insight to understand tumor progression. In the current study, the hypothesis is that evolutionary paths impact on tumor heterogeneity. Multiple evolutionary paths would lead to high tumor heterogeneity. Additionally, dominated timing of the major evolutionary path also made effect on cancer progression. To verify this point, we reconstructed pan-cancer evolution process by BML using mutation data, and identified four pan-cancer evolution patterns based on NMF clustering for 21 type of cancers: tree pattern with moderate progression, chaos pattern with high disorder, biconvex pattern with significant distinctions between early and late stages, and Cambrian pattern with an explosion in late stages. The classification based on the evolution patterns is in good accord with both clinical performance and biological evidences (e.g., gene expression and protein-protein interactions). We generated features of four evolution patterns in **Table 1**.

**Table 1:**
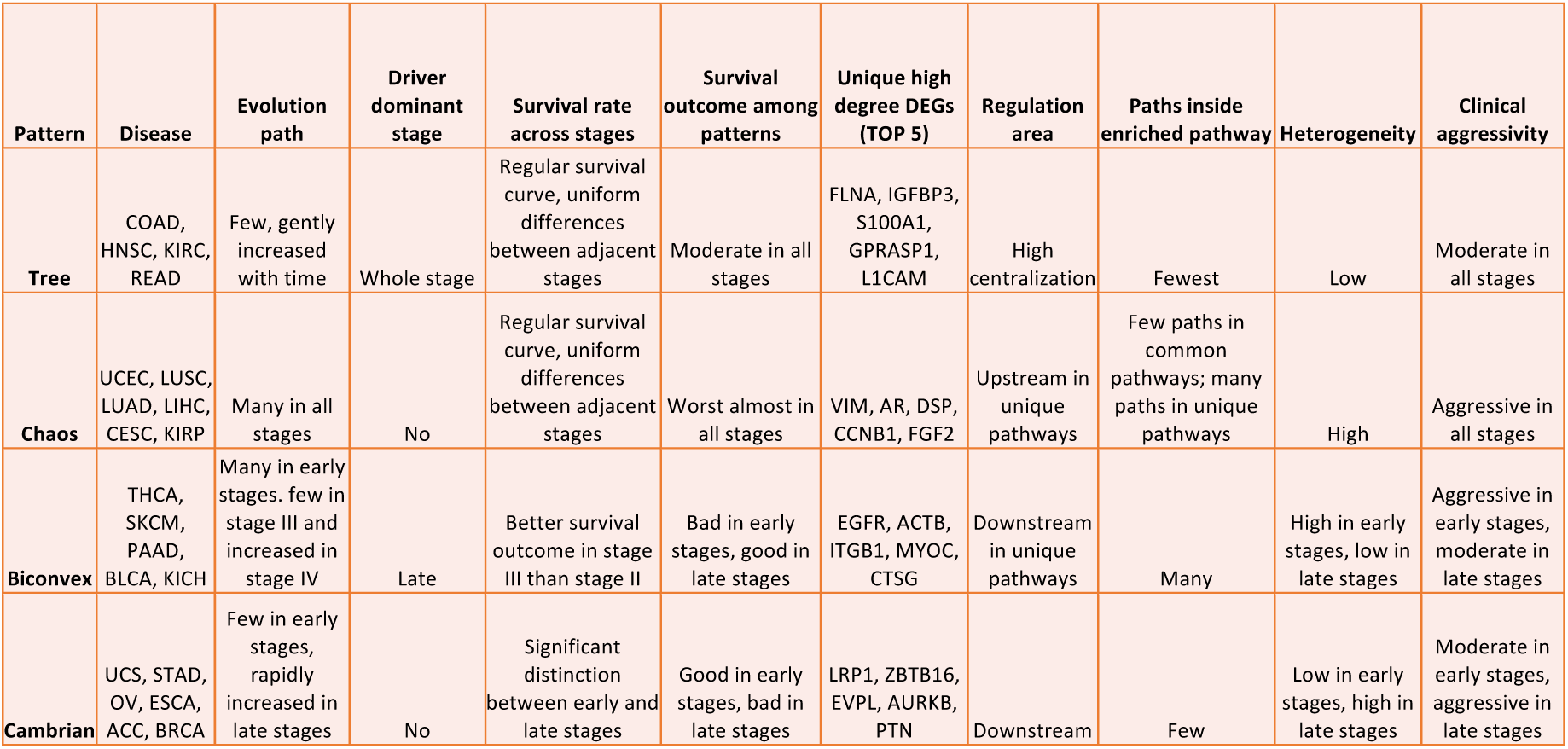
Differences of four evolution patterns. Evolution features of diseases in four evolution patterns according to our research.

Tree pattern and chaos pattern are the typical evolution patterns. The former employs a major evolutionary path, leading to a comparatively low tumor heterogeneity according to our hypothesis. Cancers in this evolution pattern, e.g., COAD and READ, also have remarkable driver genes(Sottoriva et al. 2015; Pang et al. 2018; Alexander Davis, Ruli Gao 2017). Due to the low heterogeneity and smoothly progression, tree pattern showed an optimistically clinical aggressivity. Chemical and immune Therapies targeting these driver genes in tree pattern can receive a miraculous curative effect. The latter is completely out of order, and none of the evolutionary paths showed majority. The rough-and-tumble evolutionary paths and unclear evolution progression in chaos pattern lead to high tumor heterogeneity, resulting in aggressive survival outcomes. Lung cancer is a typical example for chaos pattern, which shows a remarkable tumor heterogeneity in clinical cases(Liu et al. 2016). Biconvex pattern is a mixture of tree pattern and chaos pattern. Similar with chaos pattern, biconvex pattern exhibits a disordered feature in evolutionary paths in early stages. As no major evolutionary path or remarkable driver genes are detected, tumors in this evolutionary path have a comparatively high heterogeneity, resulting in a poor survival performance. However, after forming a dominant evolutionary paths in stage III, biconvex pattern shows a similar behavior to tree pattern, and have a better survival outcome compared to stage II. For the cancers in biconvex pattern, clinical treatment targeting stage III will receive a better efficacy(Krishnan et al. 2017). Cambrian pattern is a special one, because of having an explosion of evolutionary paths. Before explosion, this evolution pattern has a smooth tumor progression and shows a good survival performance, which suddenly drops off after explosion. This means that patients in this evolution pattern always suffer an emergency circumstance(Poveda et al. 2014). In conclusion, tree pattern showed a high order evolution process and resulted in optimistically clinical aggressivity. The high tumor heterogeneity in Chaos pattern and early-biconvex pattern drove poor survival performance. While late-biconvex pattern was better organized and reduced its clinical aggressivity. Cambrian pattern showed a good survival performance until the explosion happened, which sharply increased the clinical aggressivity of tumor.

Genes with high PPI and cancer-connection degrees, e.g., DES(Ellis et al. 2012; Seshagiri et al. 2012) and DCN(Network et al. 2011; Muzny et al. 2012), played essential roles in cancers, and their expression had significant impacts on tumor environments. MMP9, MMP2, DCN, COL1A1, SPP1 and CAV1 were experimentally confirmed key genes for cancers(Huang et al. 2016; Chai et al. 2016). The matrix metalloproteinase (MMP) family (MMP9 and MMP2) always functioned with growth factors, and were associated with inflammatory processes, indicating their critical roles in VEGF and other related hallmark pathways for cancers.

Despite various evolution paths appeared in Cambrian pattern in the late stages, their functional variation focused on minimum pathways. Most of the pathways were in downstream regulations and paths inside pathway were also limited. The explosion seemed to be an effect of system disorders accumulation. Tree pattern had the fewest paths and highest centralization regulation area, indicating throughout major evolution paths. And the biconvex pattern is consisting of early-chaos and late-tree, which coincident with its survival outcome. Compared to chaos pattern, it had more paths and downstream genes. The downstream early-chaos relieved system deterioration and resulted in better survival outcome. Additionally, in the cell adhesion molecules pathway DEGs in chaos pattern were exempted from immune system compared to biconvex pattern. The disturbance of immune system could bring out a severe evolution progression.

Our research reconstructed pan-cancer evolution process based on somatic mutations across four pathological stages. We proposed four cancer evolution patterns which is in consistent with their survival outcome. Except study based on genomic data, we also used gene expression data for functional enrichment analysis and explored their similarities and differences. On the other hand, we found some DEGs with high PPI degree and cancer-connection which should be valued. Our study therefore furthers the understanding of tumor progression and figured out how they drive clinical aggression.

The unbalance sample size and heterogeneity among different patients would be limiting factors for cancer evolution study. We used the bootstrap method to construct the evolution process and only picked out highly convincible genes (**see Methods**). The clinical aggressivity and function analysis accordant with this evolution model and advanced the understanding of tumor progression progress. On account of the different evolution patterns of different cancers, the optimum treatment time would be helpful to remit clinical aggressivity. Additionally, variations in downstream and upstream of biological pathways have distinct effects. In general, drugs targeted on upstream genes always have a better therapeutic outcome, while consideration of evolution pattern would make biomarker selection more meaningful.

## Materials and Methods

### Data Processing

All pan-cancer samples derived from TCGA Data Portal Bulk Download (http://tcga-data.nci.nih.gov/tcga)(Chang et al. 2013), with a declaration that all TCGA data are now available without restrictions on their use in publications or presentations. We used 21 kinds of cancer in total. Somatic nucleotide variants (SNV) used for the following study were subsequently annotated by Oncotator(Ramos et al. 2015) in UCSC Xena (http://xena.ucsc.edu), only those curated SNVs were picked out. SNV data summary and cancer descriptions are generated in **Table S1**. Cancer detection time and the biological system were obtained from (http://www.cancer.org). And M/C class annotation was derived from Ciriello’s article(Ciriello et al. 2013). Patients have extinct pathological stage clinical information were kept while others were filtered. After removing hypermutated samples and genes with low mutation frequency (<3), we transformed them into a 0/1 matrix (patient x mutation gene). The correlation heatmap (**Fig. 1d**) was performed by hierarchical cluster using the median and mean of gene mutation frequency and 5-year survival rate. The 5-year survival rate for each cancer was calculated using TCGA dataset, and we also evaluated cancer prognosis by existing research (http://www.cancersurvivalrates.net).

### Reconstruction of pan-cancer evolution process

Cancer evolution process was reconstructed using the approach we published before(Pang et al. 2018). Combining with probability network reconstructed by Bayesian mutation landscape (BML)(Misra et al. 2014), we generated evolutionary paths including genes with both high and moderate mutation frequency. After built DAG map using raw data (**Fig. S1**), we generated convincible evolution paths using bootstrap score threshold. We randomly selected 30 samples (with replacement) at each stage for 100 times in case sample bias. Nodes appeared more than 60 times, and nodes appeared more than 10 times in each pathological stage of particular cancer were kept. Genes appeared more than once in combined and separate pathological stages DAGs in the raw map were recognized as DAG key genes. These three vectors of four pathological stage were used for NMF cluster. An R script implemented this clustering process by R package “NMF”(Gaujoux and Seoighe 2010). Four evolution pattern figures (**Fig. 2c**) were manually sketched. Evolutionary paths with direct connection to normal node and had more than one key genes was considered as major evolutionary path.

### Survival analysis of cancers in the same pattern

Survival time used in this paper was the time to death or censor event. Survival curve in **Fig. 3** was generated by Kaplan-Meier estimator and plotted by R package “survminer”. Survival analysis in **Table S3** was performed using R package “survival”(Harrington and Fleming 1982).

### Protein-protein interaction network of differentially expressed gene and functional enrichment analysis

Tumor gene expression data were obtained from TCGA, too. Since we only wanted to find different expression genes rather than precise quantify, gene expression data were not matched with SNV data. We used GEPIA database(Tang et al. 2017) as supplements for cancers without gene expression data in TCGA. After construct disease-gene network, we added protein-protein interaction from Human Protein Reference Database (HPRD, http://www.hprd.org) (Keshava Prasad et al. 2009). Network construction and analysis were generated by Cytoscape(Shannon et al. 2003). High disease-connected DEGs and high PPI degree DEGs were collected in **Table S5**. We picked out hub DEGs (degree>5) for functional enrichment using WEB-based GEne SeT Analysis Toolkit(Wang et al. 2013) for with parameters set as Bonferroni, p<0.05.

After that, we separated enriched KEGG pathways to two parts, and defined genes with more downstream regulations than upstream as upper genes. Under genes referred to the opposite.

Regulation area was related to the amount of upper and under genes.

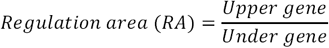

We also counted paths in individual KEGG pathways, and genes with direct or indirect connection (irreversible direction) were supposed to be in the same path. RA normalization was performed global and path normalization was performed among patterns.

## Acknowledgements

This work was supported by the National Key R&D Program of China (2016YFC0901704, 2017YFA0505500, 2016YFC0902400) and the Youth Innovation Promotion Association CAS (2017325).

## Author contributions

S.C.P designed the evolution reconstruction process and carried out the analysis. X.S helped with the algorithm. S.C.P, Y.D.S and L.L.W prepared Figures. S.C.P and J.F.W. wrote the main manuscript text. Z.W, Y.L.Z and Y.X.L conceived and supervised the experiments. All authors reviewed the manuscript.

## Additional Information

### Competing interests

The authors declare no competing financial interests.

**Corresponding author**

**Correspondence to:** YiXue Li, Zhen Wang, Yi-Lei Zhao and Jingfang Wang

